# Accelerated protein retention expansion microscopy using microwave radiation

**DOI:** 10.1101/2024.05.11.593228

**Authors:** Meghan R. Bullard, Juan Carlos Martinez Cervantes, Norisha B. Quaicoe, Amanda Jin, Danya A. Adams, Jessica M. Lin, Elena Iliadis, Tess M. Seidler, Isaac Cervantes-Sandoval, Hai-yan He

## Abstract

Protein retention expansion microscopy (ExM) retains genetically encoded fluorescent proteins or antibody-conjugated fluorescent probes in fixed tissue and isotropically expands the tissue through a swellable polymer network to allow nanoscale (<70 nm) resolution on diffraction-limited confocal microscopes. Despite numerous advantages ExM brings to biological studies, the full protocol is time-consuming and can take multiple days to complete. Here, we adapted the ExM protocol to the vibratome-sectioned brain tissue of *Xenopus laevis* tadpoles and implemented a microwave-assisted protocol to reduce the workflow from days to hours. In addition to the significantly accelerated processing time, our microwave-assisted ExM (^M/W^ExM) protocol maintains the superior resolution and signal-to-noise ratio of the original ExM protocol. Furthermore, the ^M/W^ExM protocol yields higher magnitude of expansion, suggesting that in addition to accelerating the process through increased diffusion rate of reagents, microwave radiation may also facilitate the expansion process. To demonstrate the applicability of this method to other specimens and protocols, we adapted the microwave-accelerated protocol to whole mount adult brain tissue of *Drosophila melanogaster* fruit flies, and successfully reduced the total processing time of a widely-used *Drosophila* IHC-ExM protocol from 6 days to 2 days. Our results demonstrate that with appropriate adjustment of the microwave parameters (wattage, pulse duration, interval, and number of cycles), this protocol can be readily adapted to different model organisms and tissue types to greatly increase the efficiency of ExM experiments.

## 1. Introduction

Microscopy is an indispensable tool in the study of cellular and subcellular structures and functions. The advent of optical fluorescence microscopy marked a new era of the biological sciences as it not only allows for selective detection and localization of specific molecules and proteins in various biological tissues, but also provides a large toolkit of various fluorescent probes for simultaneous multicolor (Bayguinov et al 2018, Yuste 2005). Despite these advantages, the resolution of optical microscopy is limited to a few hundred nanometers due to the optical diffraction limit (Sezgin 2017). Expansion microscopy is a recently developed technique that uses a swellable polymer network to physically expand the tissue in order to overcome the diffraction limit of conventional optical microscopy and achieve higher resolution (Chen et al 2015). Protein retention expansion microscopy (proExM, abbreviated hereafter as ExM for simplicity) is one type of expansion microscopy that uses a small molecule cross-linker, succinimidyl ester of 6-((acryloyl)amino) hexanoic acid (AcX), which modifies the amine groups on proteins to facilitate the anchoring of genetically encoded fluorescent proteins or antibody-conjugated fluorophores to a swellable gel and preserves the fluorescence for subsequent imaging following expansion (Tillberg et al 2016). This method allows for the visualization of fluorescently labeled proteins of interest in fixed biological specimens using diffraction-limited microscopes with the resolution at tens of nanometers (Campbell et al 2021, Cheng et al 2023, Martinez et al 2020, Sheard et al 2019, Sneve & Piatkevich 2021, Tillberg et al 2016).

Despite the superior resolution and other advantages such as easy accessibility and reduced light scattering through the optical clearing of the tissue, the complete ExM protocol takes 2-3 days with multiple cumbersome incubation steps (Asano et al 2018). This is independent of immunohistochemistry (IHC) which can increase the protocol to over seven days depending on the incubation duration required by the antibodies. Although many efforts have been made to optimize ExM for various applications, none of the efforts have addressed the prolonged incubation time. Here we present an accelerated protocol that combines ExM with microwave radiation to accelerate the process without compromising the immunostaining quality and expansion efficiency. Microwave radiation has long been used to successfully accelerate and enhance tissue fixation for immunofluorescence microscopy and electron microscopy (Bond & Kinnamon 2013, Ferris et al 2009, Giberson et al 2003, Leong & Milios 1990, Miura et al 1988, Munoz et al 2004). It can also facilitate and accelerate the primary and secondary antibody incubation steps in immunostaining protocols in multiple tissue types (Bond & Kinnamon 2013, Chiu & Chan 1987, Ferris et al 2009, Munoz et al 2004). The acceleration process likely works through both thermal and non-thermal effects that facilitate the diffusion process (Galinada & Guiochon 2007, Hinrikus et al 2015). One potential caveat of using microwave to process biological tissue is uncontrolled excessive heating inside the tissue, creating uneven hot spots which can compromise the tissue integrity as well as fine subcellular structures. This can be avoided with the utilization of a specially designed microwave with automated temperature control, which cycles tissue specimens in working solutions through on and off cycles in a precise temperature-controlled environment with user-defined parameters (Munoz et al 2004).

Here we report an accelerated ExM protocol using microwave radiation (^M/W^ExM) which we adapted from a previously published ExM protocol (Asano et al 2018) and microwave accelerated IHC protocols (Ferris et al 2009, Munoz et al 2004) (^M/W^IHC; Fig. 1A-B). We first tested various parameters and optimized the ^M/W^IHC and ^M/W^ExM for single and double immuno-labeling in vibratome sections of fixed brain tissue of *Xenopus laevis* tadpoles using a commercially available specialized microwave with precise temperature control. To demonstrate the versatility of our ^M/W^ExM protocol, we then adapted these protocols to the brain tissue of *Drosophila melanogaster* and tested the applicability of using microwave-assisted processing to accelerate the IHC-ExM protocols in the whole mount *Drosophila* brain. The optimized ^M/W^IHC-^M/W^ExM protocol of the fruit fly whole mount brain reduced the processing times by at least 15-fold compared to standard IHC-ExM protocols, greatly increasing the efficacy of the experimental protocol.

**Figure 1.**
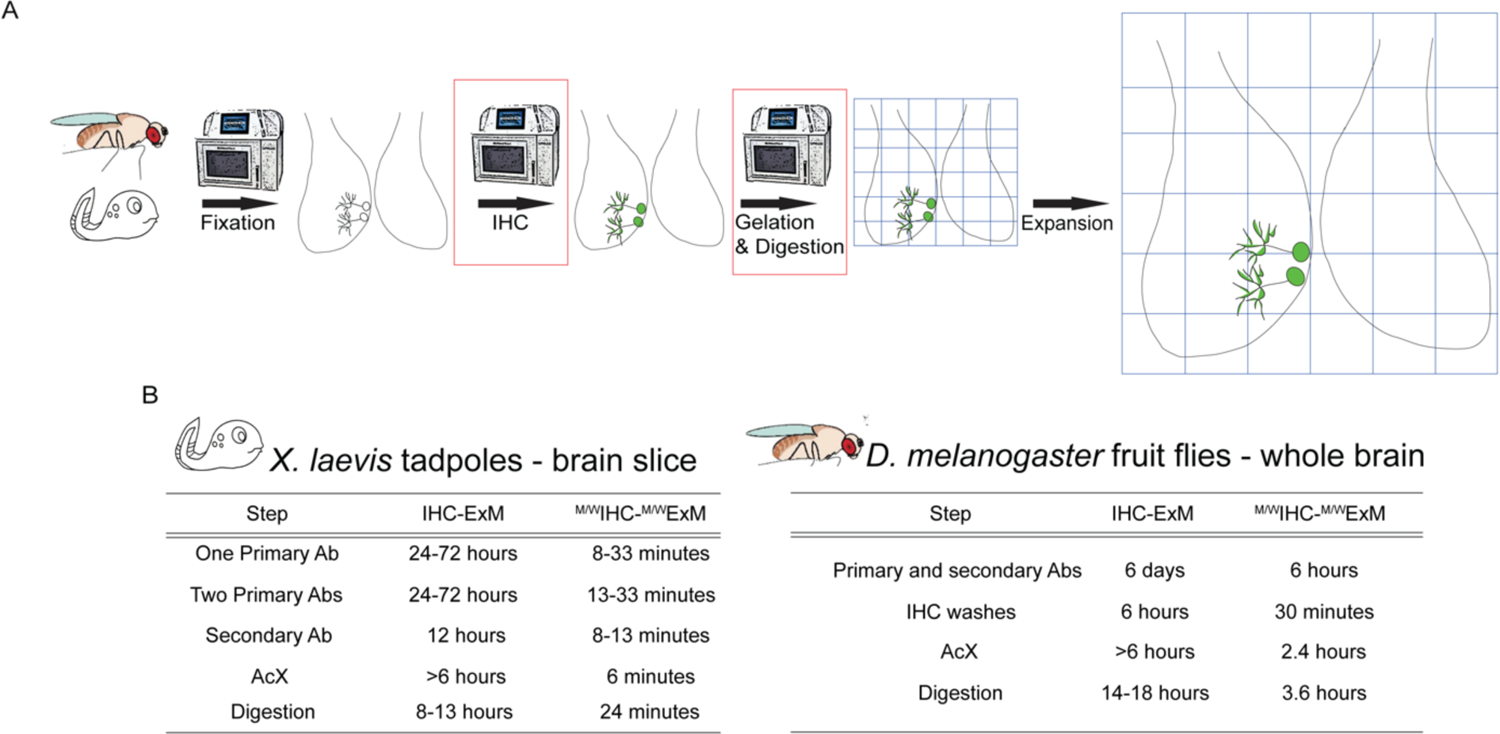
Workflow for microwave assisted accelerated ExM. A. Timeline of ExM in *Xenopus laevis* and *Drosophila melanogaster* beginning with fixation. Steps boxed in red are accelerated with microwave radiation. B. Table comparing incubation times for accelerated steps between conventional and microwave-assisted protocols.

**Figure 2.**
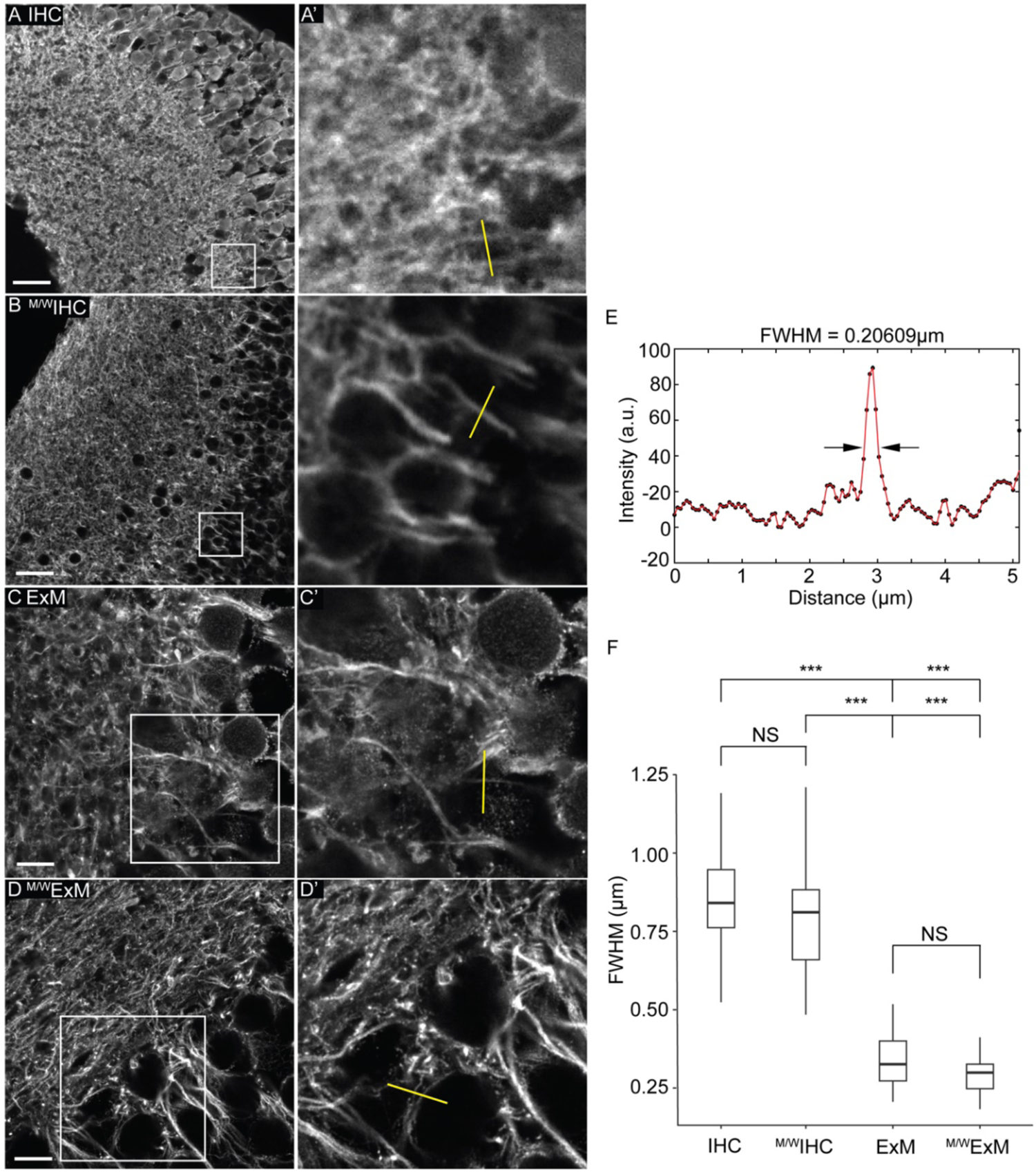
Microwave radiation accelerates IHC-ExM in the *Xenopus laevis* brain tissue. A-D. Representative confocal images of vibratome sections of optic tectum immunostained with anti-β-tubulin antibodies. A’-D’. Magnification of boxed regions in (A-D). Yellow line depicts examples of 5μm line ROIs drawn for FWHM analysis. A-B. Non-expanded sections immunostained using conventional (IHC, A) or microwave accelerated (^M/W^IHC, B) protocols. Scale bars: 25μm. C-D. Expanded sections processed with conventional (ExM, C) or microwave accelerated ExM (^M/W^ExM, D) protocols. Scale bars: 6.25μm. E. Representative intensity plot of a line ROI from an ExM image (scale calibrated by linear expansion factor). Solid red line shows the line fit used to determine the full-width at half-maximum (FWHM). F. Summary data for FWHM. Whiskers depict interquartile range. Black line depicts median. Same in the rest of figures. ***: *p <* 0.0001, Kruskal-Wallis rank sum test with Dunn’s Test for multiple comparisons.

Both *Xenopus laevis* tadpoles and *Drosophila melanogaster* fruit flies are important model organisms for biological research. *Xenopus laevis* has been used for almost a century for biological studies and has been instrumental in developmental and neuroscience research (Dawid & Sargent 1988, Exner & Willsey 2021, Gurdon & Hopwood 2000, Lee-Liu et al 2017, Straka & Simmers 2012). The development of their neural circuit has been studied extensively, especially the early stages of embryonic development such as neural induction and patterning (Exner & Willsey 2021). The visual circuit of *Xenopus laevis* tadpoles is a well-established model system for visually driven plasticity paradigms of vertebrate neural circuits and has contributed significantly to our understanding of activity-dependent plasticity mechanisms (Cline et al 2023, Cohen-Cory et al 2010, Dingwell et al 2000, Liu et al 2016, Richards et al 2010, Ruthazer & Aizenman 2010). More recently, *Xenopus* tadpoles have also begun to be used as disease models for neurodegenerative disorders such as Alzheimer’s, Parkinson’s, and Huntington’s Disease as well as other disorders such as multiple sclerosis and Fragile X syndrome, among others (Faulkner et al 2015, Haremaki et al 2015, Horowitz et al 2001, Kaya et al 2012, Liu et al 2018, Mannioui et al 2018, Paganelli et al 2001, Pratt & Khakhalin 2013, Sater & Moody 2017, Truszkowski et al 2016, Willsey et al 2020). Here we show for the first time that ExM can be adapted to visualize subcellular components in *Xenopus laevis* neural tissue and used this system to test and demonstrate the feasibility of using microwave-assisted processing to accelerate the ExM protocol. Our microwave-assisted ExM (^M/W^ExM) protocol significantly shortens the processing time for IHC-ExM experiment and maintains the superior resolution and signal-to-noise ratio of the original ExM protocol. Furthermore, the ^M/W^ExM protocol consistently yields higher expansion, suggesting that microwave radiation may also facilitate the expansion process.

*Drosophila melanogaster* is another organism that has been widely used in biological research for over a century to model complex biological questions. They are easy and inexpensive to culture in laboratory conditions, have a relatively short life cycle, and produce large numbers of externally laid embryos (Hales et al 2015). It is a powerful model organism with great amenability to genetic manipulations. There also exists a rich repertoire of open-source databases like FlyBase (Gramates et al 2017) and the full brain connectome (Winding et al 2023). Neurobiologists have used these tools to investigate brain development and physiology at different spatiotemporal scales and have provided mechanistic insights into brain function (Sjulson et al 2016). Our adaptation of the ^M/W^IHC-^M/W^ExM protocol to the whole brain fly tissue significantly improved the efficiency of the current working protocol. By completing the most time-consuming steps in the microwave, we successfully reduced the total processing time of a widely used *Drosophila* IHC-ExM protocol (Asano et al 2018, Jenett et al 2012) from 6 days (excluding the fixation time) down to 2 days (Fig.1B). We confirmed that both ^M/W^IHC and ^M/W^ExM preserve tissue integrity and maintain the gross cellular morphology. Consistent with our observation in the *Xenopus* tissue, we also found that tissue expanded with the ^M/W^ExM protocol exhibited higher linear expansion than conventional ExM method in *Drosophila* tissues. In summary, we demonstrated here the microwave-assisted processing significantly increases the efficiency of the ExM protocol with increased linear expansion while maintaining the superior resolution of the conventional ExM method. With appropriate modifications and optimizations, this protocol has the potential to be adapted to other types of ExM methods as well as other organisms and tissue types.

## 2. Material and Methods

### 2.1 *Xenopus laevis* husbandry and electroporation

All animal protocols were approved by the Georgetown University Institutional Animal Care and Use Committee. Albino *Xenopus laevis* tadpoles were either fertilized in-house or purchased from Xen Express (Brooksville, Florida). Tadpoles were reared at 22–23 °C with 12h dark/12h light cycle in 0.1x Steinberg’s solution (58.0mM NaCl, 0.67mM KCl, 0.34mM Ca(NO_3_)2.4H_2_O, 0.83mM MgSO4.7H_2_O, 0.5mM Tris, pH 7.5). Animals began to be fed at stage 47 (J 1994, McKeown & Cline 2019). Stage 47-49 animals were used for all experiments. For electroporation, animals were anesthetized in 0.02% MS-222 (Tricane methanesulfonate, Sigma, St. Louis, MO), and the tectum was co-electroporated with CMV::Gal4 (2µg/µL) and UAS-synaptophysin-GFP (5µg/µL), at 35V and 3-4 pulses with 1.6msec pulse duration, as previously described (He et al 2018). All animals were screened 3-6 days post-electroporation for proper level of the synaptophysin-GFP expression in tectal neurons before fixation.

### 2.2 Drosophila melanogaster husbandry

Flies were cultured on standard food at room temperature. Flies used for the endogenous expression pattern of Scribble protein correspond to a protein trap line, Scrb::GFP^CA07683^. For the rest of the experiments, *uas-dcr2/+; nsyb-gal4* flies with a CantonS genetic background were used.

### 2.3 Immunostaining of *Xenopus laevis* free-floating vibratome sections

On the day of fixation, animals were anesthetized in 0.02% MS-222 and then immersed in 4% PFA (diluted from 16% PFA in 1xPBS, Cat # 15710, Electron Microscopy Sciences, Fort Washington, PA) using a Pelco Biowave Pro microwave (Model 36700, Ted Pella, Redding, CA. 1-3 animals/well, 1mL fixative/ well in a 24-well plate, 150W 1 min on, 1 min off, 1 min on) followed by a post-fix at 4° C overnight.

Animals were then washed in 1x PBS for 5 minutes and then brains were dissected. Dissected brains were embedded in 20% chicken albumin (Sigma-Aldrich, catalog #A5253, St. Louis, MO) with 1.5% gelatin; (Sigma-Aldrich-Aldrich, catalog #G1890, St. Louis, MO) made in double distilled H_2_O (ddH_2_O) and crosslinked by 1% glutaraldehyde (diluted from 50% glutaraldehyde, Electron Microscopy Sciences, catalog #16320, Hatfield, PA). 40 µm sections were cut on a vibratome (Leica VT1000S) for free-floating section immunostaining. Sections were incubated in 1% sodium borohydride (Sigma-Aldrich, catalog #AC189300000, St. Louis, MO) in 1% PBSTw (1% Tween-20 in 1xPBS) for 15 min to quench autofluorescence and then washed in 1xPBS for 3×10 min. Sections were then blocked in 10% normal goat serum (NGS; Thermo Fischer Scientific, catalog #16201, Waltham, MA) and 1% fish gelatin (Thermo Fischer Scientific, catalog #40600036, Waltham, MA) in 1% PBSTw for 1 hour in room temperature.

After blocking, sections were incubated in primary antibody (mouse anti-β-tubulin I&II, Sigma-Aldrich; catalog #T8535, RRID: AB_261795, 1:500 in 1% normal goat serum and 1% fish gelatin in 1% PBSTw) for 24 hours in 4°C on a rotator. After 3 x 15 min rinses with 1% PBSTw, sections were incubated in secondary antibodies (Goat anti-mouse CF633; Sigma-Aldrich; catalog #SAB4600139; 1:1000) diluted in 1% normal goat serum and 1% fish gelatin in 1% PBSTw overnight in 4° C. After 3 x 10 min washes in 1xPBS, sections were either mounted on slides in Vectashield Hardset Mounting Media (Vector Laboratories, catalog #NC9029228, Burlingame, CA) and hardened in room temperature in dark for same-day imaging or expanded using conventional expansion protocol described below (section 2.5).

### 2.4 Immunostaining of *Drosophila* whole brains

Fly whole brains were dissected and immunolabeled as previously described (Jenett et al 2012) with some modifications. Briefly, male and female flies were anesthetized on ice or with carbon dioxide gas. Flies were first rinsed once in 70% ethanol, then washed in ice-cold S2 media (Schneider’s Insect Medium, Sigma) on a dish well and dissected in a second well containing the same cold media. After dissection, each brain was immediately transferred to a 1.7 mL Eppendorf LoBind microcentrifuge tube containing 1 mL of 1% paraformaldehyde in S2 media on ice. A tube rotator at a fixed speed (24 rpm) was used to mix overnight at 4 °C. After 16-24 h, the fixative solution was removed by aspiration and replaced immediately with 1 mL of ice-cold PAT (0.5% Triton X-100, 0.5% bovine serum albumin in 1X PBS) and mixed by hand inverting a few times. The liquid was then discarded and replaced with two more washes of PAT, each wash at room temperature for 1 h. For blocking, brains were incubated in 1 mL of 3% normal goat serum (BH) in PAT for 1.5 h at room temperature. Brains were then incubated in 1 mL of primary antibody solution in blocking buffer (1:50 monoclonal mouse anti-nc82, Developmental Studies Hybridoma Bank, Iowa City, IA; or 1:500 rabbit anti-GFP, Thermofisher cat# A11122) at room temperature for 3 h in constant rotation. Then, the samples were moved to 4 °C overnight. The next day, the primary antibody solution was aspirated and immediately replaced with 1 mL of ice-cold PAT and mixed by hand for a few seconds. The liquid was then discarded and replaced with two more washes of PAT, each wash at room temperature for 1 h. The secondary antibody solution was prepared in blocking buffer (CF488A or CF633A anti-mouse or anti-rabbit antibodies, Biotium cat# 20020 and 20124, 1:800). Once again, samples were incubated at room temperature with constant rotation for 3 h followed by an incubation at 4 °C for 5 days. After secondary antibody incubation, three washes with PAT were performed as described above followed by a final wash with 1X PBS. This was the last step before mounting on a glass slide with Vectashield.

For SYTOX Green staining, brains were fixed overnight, followed by three washes as indicated above. An additional wash was done with 1X PBS. The nuclei were stained by 20 min incubation in 1:3000 SYTOX Green in 1X PBS at room temperature without agitation. After incubation, two immediate washes with 1X PBS were performed. Brains were finally mounted in Vectashield.

### 2.5 Microwave accelerated immunostaining of *Xenopus laevis* free-floating vibratome sections

40 µm free-floating vibratome sections were prepared as described above (see section 2.3). Various primary antibody incubations require differing numbers of 3 min on - 2 min off cycles (Table 1). We found that optimal protocols range between 2∼7 cycles of the 150 W 3 min on - 2 min off cycle (Table 1), depending on the antibodies used as well as the number of antibodies used for co-immunostaining (see results for details). Alternatively, when used in conjunction with ExM, the protocol can be spread out into a two-day protocol for more convenient experimental timing by incubating secondary antibodies overnight at 4° C on rotator (see section 2.3). The washes following both primary and secondary antibody labeling were done exactly as described in the conventional protocol. Similarly, non-expanded sections were mounted as described above and hardened at room temperature for same-day imaging. Sections that were not mounted were processed using ^M/W^ExM (see section 2.7).

**Table 1.**
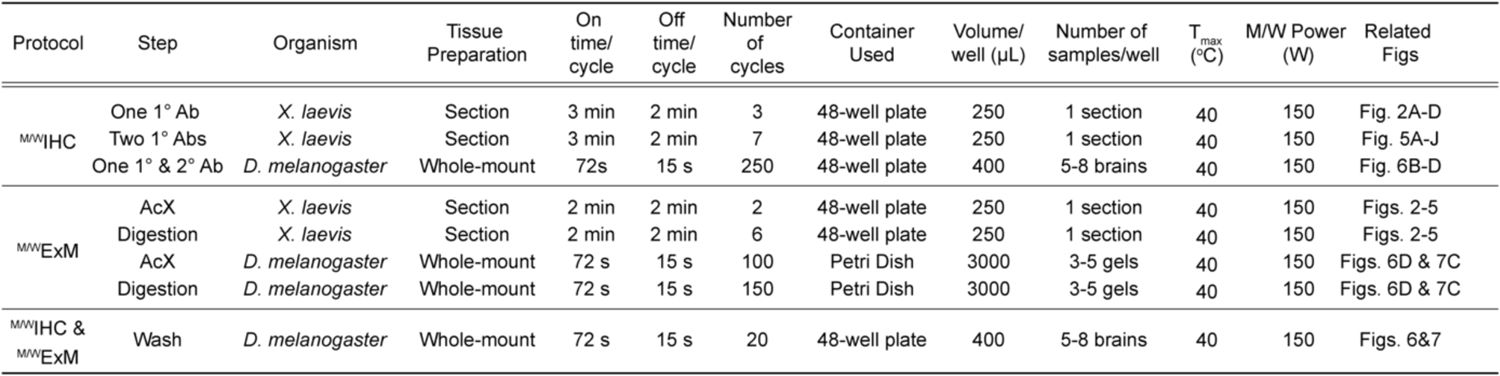
Key parameters for microwave-processing steps used.

| Protocol | Step | Organism | Tissue Preparation | On time/ cycle | Off time/ cycle | Number of cycles | Container Used | Volume/ well (μL) | Number of samples/well | T<sub>max</sub> (°C) | M/W Power (W) | Related Figs |
| --- | --- | --- | --- | --- | --- | --- | --- | --- | --- | --- | --- | --- |
| <sup>M/W</sup>IHC | One 1° Ab | *X. laevis* | Section | 3 min | 2 min | 3 | 48-well plate | 250 | 1 section | 40 | 150 | Fig. 2A-D |
|  | Two 1° Abs | *X. laevis* | Section | 3 min | 2 min | 7 | 48-well plate | 250 | 1 section | 40 | 150 | Fig. 5A-J |
|  | One 1° & 2° Ab | *D. melanogaster* | Whole-mount | 72 s | 15 s | 250 | 48-well plate | 400 | 5-8 brains | 40 | 150 | Fig. 6B-D |
| <sup>M/W</sup>ExM | AcX | *X. laevis* | Section | 2 min | 2 min | 2 | 48-well plate | 250 | 1 section | 40 | 150 | Figs. 2-5 |
|  | Digestion | *X. laevis* | Section | 2 min | 2 min | 6 | 48-well plate | 250 | 1 section | 40 | 150 | Figs. 2-5 |
|  | AcX | *D. melanogaster* | Whole-mount | 72 s | 15 s | 100 | Petri Dish | 3000 | 3-5 gels | 40 | 150 | Figs. 6D & 7C |
|  | Digestion | *D. melanogaster* | Whole-mount | 72 s | 15 s | 150 | Petri Dish | 3000 | 3-5 gels | 40 | 150 | Figs. 6D & 7C |
| <sup>M/W</sup>IHC & <sup>M/W</sup>ExM | Wash | *D. melanogaster* | Whole-mount | 72 s | 15 s | 20 | 48-well plate | 400 | 5-8 brains | 40 | 150 | Figs. 6&7 |

### 2.6 Microwave accelerated immunostaining of whole-mount *Drosophila melanogaster* brains

Fly whole brains were dissected and fixed as described in 2.4. The *Drosophila* microwave cycle alternates with 72 s *on* and 15 s *off*. This cycle is repeated for various numbers for each of the steps described below (Table 1). For all the IHC steps performed in the microwave, dissected brains were transferred into a 48-well plate.

Each well contained 5-8 brains in a working volume of 400 μL/ well. To avoid evaporation of the solutions during the microwave cycles, the well plate was covered with the lid. The temperature probe was immersed in a separate well of the same plate containing 400μl 1X PBS. A hole was drilled in the lid to ensure the access of the probe to the well. After fixation, the fixative solution was replaced with PAT. Two more washes with PAT were done in the microwave with 20 cycles each time (Table 1). The blocking step was done as described in section 2.4. Different from the conventional IHC protocol described in section 2.4, instead of sequential incubation in primary and secondary antibodies, the samples were then incubated in both the primary and secondary antibodies simultaneously for 250 microwave cycles (Table 1). After the antibody incubation step, the samples were washed three times in PAT with 20 microwave cycles each time. A final 1X PBS wash was performed before either mounting the brains or proceeding to ExM.

### 2.7 Gelation, digestion, and expansion of immuno-stained vibratome sections of tadpole brain tissue

Protocols in this section were adapted from Asano et al., 2018. In order to preserve the fluorescence signal in the immuno-stained tissue sections, all ExM steps were performed in foil-wrapped containers to minimize exposure to light. Stock AcX solution (10mg/mL) was prepared by dissolving AcX (ThermoFisher, catalog # A20770, Waltham, MA) in DMSO and stored at −20 °C. Sections were incubated in AcX solution diluted 1:100 in 1x PBS for at least 6 hours at room temperature. Sections were then washed 2 x 15 min in 1xPBS. A gelation solution was prepared with Stock X (g/100mL 1xPBS: sodium acrylate 8.6, acrymalide 2.5, N,N’-methylenebisacrylamide 0.15, NaCl 11.7), 4-Hydroxy TEMPO (4HT, Sigma-Aldrich, catalog #176141, St. Louis, MO, 0.5% wt/vol in H_2_O), N’,N’,N’,N’-tetramethylethylenediamine (TEMED; Sigma-Aldrich, catalog #T9281, St. Louis, MO, 10% wt/vol in H_2_O) and ammonium persulfate (APS; Sigma-Aldrich, catalog #A3678, St. Louis, MO, 10% wt/vol in H_2_O) in a 47:1:1:1 ratio and put back on ice to prevent premature polymerization. Tissue sections were mounted on gelation chambers made from poly-l-lysine coated glass slides and then incubated in gelation solution on ice for 30 minutes. Coating gelation chambers with poly-l-lysine helps to anchor the tissue section to the glass slide, which ensures that the section remains flat in the gel. The anchoring also ensures that the tissue section is accessible for high-NA low-working distance objectives for imaging following expansion. Poly-l-lysine coated chambers and coverslips were prepared by washing glass microscope slides and glass coverslips with acid alcohol (1% HCl in 70% ethanol) and placing 1:10 0.1% poly-l-lysine (Electron Microscopy Sciences, catalog #19320, Hatfield, PA) in ddH20 on slide for 1 hour before aspirating off and leaving covered in fume hood overnight (Suvarna SK 2013, V. 2019). Coated slides can be stored at 4° C. The tissue section covered with gelation solution in the gelation chamber was incubated at 37°C for 2 hours for the gel to polymerize. Following incubation, the cured gel was trimmed to remove the excess gel around the tissue section and incubated in proteinase K (Sigma-Aldrich, catalog #03115887001, St. Louis, MO) diluted 1:100 to 8 units/mL in digestion buffer (for 100mL H_2_O solution: 400 μL of 0.5M EDTA pH 8, 4.67g NaCl, 5mL of 1M Tris aqueous solution pH 8, 0.5g Triton X-100) for 8-13 hours in room temperature in dark.

Gels were then moved to a poly-l-lysine coated cover glass-bottom petri dish (Electron Microscopy Sciences, catalog #70673, Hatfield, PA) with a paintbrush and incubated for expansion in ddH_2_O for 3∼4 rounds of 20-minute expansion session until expansion plateaued (around 60-100 minutes). The nucleic acid stain SYTOX Green (ThermoFischer Scientific, catalog #S7020, Waltham, MA, 1:3000) can be added to the first 20-minute expansion session to stain nuclei.

### 2.8 Gelation, digestion, and expansion of immuno-stained whole *Drosophila* brains

The *Drosophila* expansion protocol was adapted from Asano *et al*. (2018). Gelation of the immuno-stained whole *Drosophila* brains was completed as described above (see section 2.7) with the following modifications: the immuno-stained brain samples were immersed in ACX solution and incubated in the gelling solution (500 μL per 6 to 10 brains) at 4°C for 30 min in the dark in a Lo Bind Eppendorf Tube. To prevent excess shaking during transfers, the gelation chamber slides were placed on a cold, flat, sturdy, transferrable surface to take the samples to the incubator later. A 200 μL pipette tip was modified by cutting approximately 3 mm of its narrow end to transfer the *Drosophila* brains onto the chamber. At this point, the brains should be oriented properly and without wrinkles. The samples were then carefully embedded in the gelling solution. To immerse the samples in the gelling solution, 20-30 μL of it were carefully added around the brains. The coverslip was then placed on top of the samples by resting on the spacers. Additional gelling solution was added to the point where the chamber was filled and without bubbles. The samples were incubated at 37 °C for at least 2 h for polymerization (we found 3 h to be optimal). After polymerization, the gel was trimmed off with a scalpel around each brain. The cured gels were then digested in ProK (diluted in digestion buffer at 1:100 to a final concentration of 8 unit/m) at room temperature in the dark for 14-16 h. To stop the digestion reaction, the digestion buffer was replaced with 1X PBS. The gels were then expanded in ddH_2_O in the dark for three rounds (20 min each round). When nucleic acids were stained, the first ddH_2_O expansion round was replaced by SYTOX Green in ddH_2_O (1:3000).

### 2.9 Microwave accelerated gelation, digestion, and expansion of immuno-stained vibratome sections and whole brains

Solutions were prepared as described above (section 2.7). The most time-consuming AcX anchoring treatment and proteinase K digestion steps were optimized in the BioWave with 100 cycles for the AcX incubation step and 150 cycles for the digestion step (Table 1). The two 1xPBS washes following the AcX incubation step were also performed in the microwave for 20 cycles each (Table 1). The rest of the steps in the protocol were performed exactly as described above (section 2.7).

### 2.10 Image acquisition, processing and data analysis

To quantify macroexpansion factor for pre- and post-expansion, gels were imaged with an Olympus MVX10 epifluorescence microscope with a MV PLAPO 0.63x objective (Fig. 4A). Confocal dual-channel imaging and SYTOX Green labeling of pre- and post-expansion *Xenopus laevis* vibratome sections was performed on an inverted LSM 880 Carl Zeiss system with a Plan-Apochromat 20x/0.8 M27 air objective (Fig. 3A- D, Fig. 4C, Fig. 5A-F). Images in the rest of the figures presented were acquired using an upright Leica TCS SP8 II system with an HC APO L20x/1.00 W water objective.

**Figure 3.**
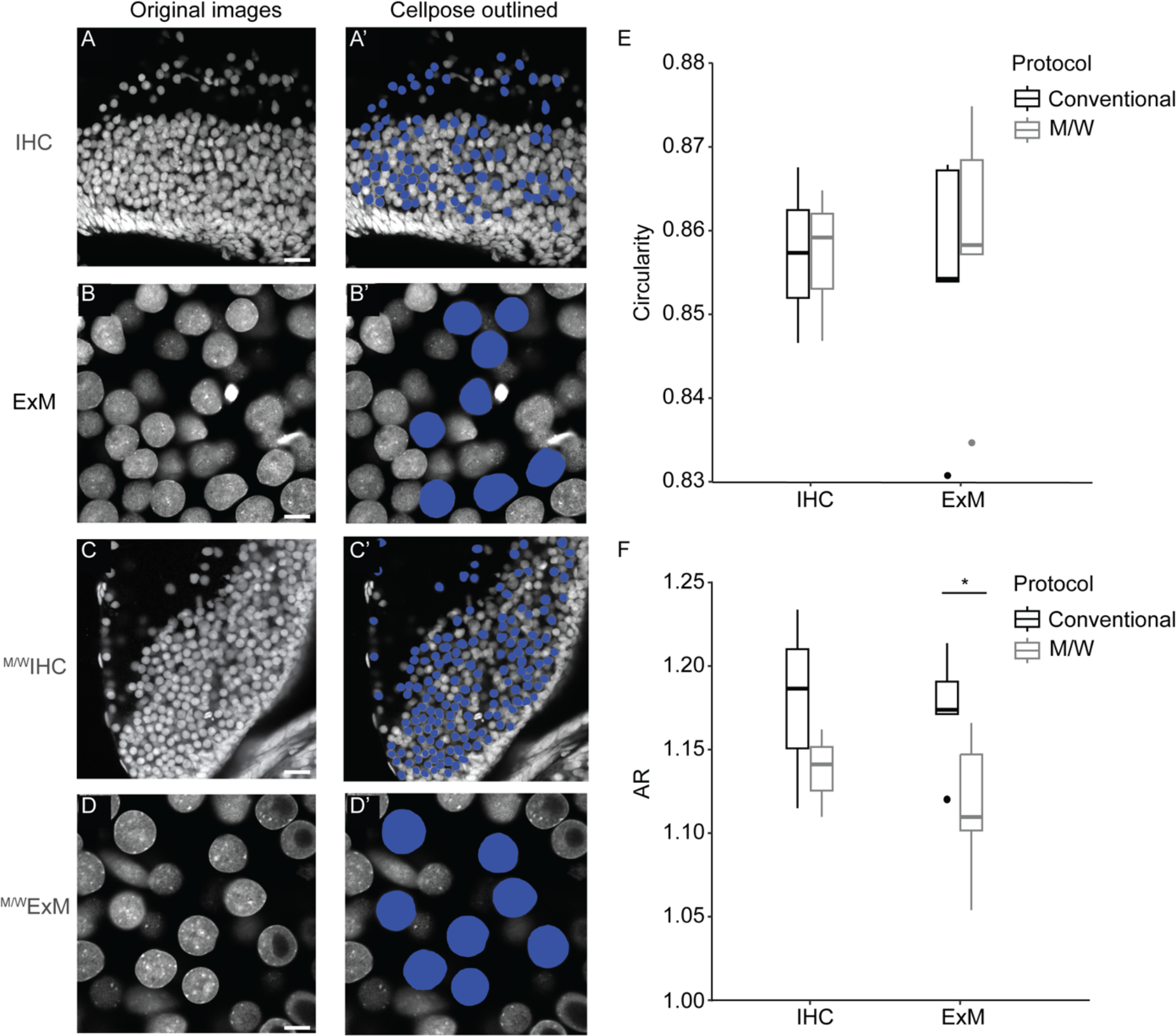
^M/W^ExM preserves gross nuclei morphology. A-D. Representative images of cell nuclei in vibratome sections of the optic tectum. A’-D’. Representative images of labelled nuclei in the optic tectum determined with the neural network Cellpose program (Stringer et al., 2021) for nuclei morphology measurement. Scale bar: 20μm (IHC & ^M/W^IHC); 5.7μm (ExM), 4.2μm (^M/W^ExM), scale bars for ExM images were calibrated by the linear microexpansion factor to match the non-expanded images. E-F. Summary data for circularity (E) and aspect ratio (AR, F) measurements of nuclei morphology. * *p* <0.05, two-way ANOVA (p = 0.029) with Fisher’s unprotected LSD post-hoc (p = 0.047).

**Figure 4.**
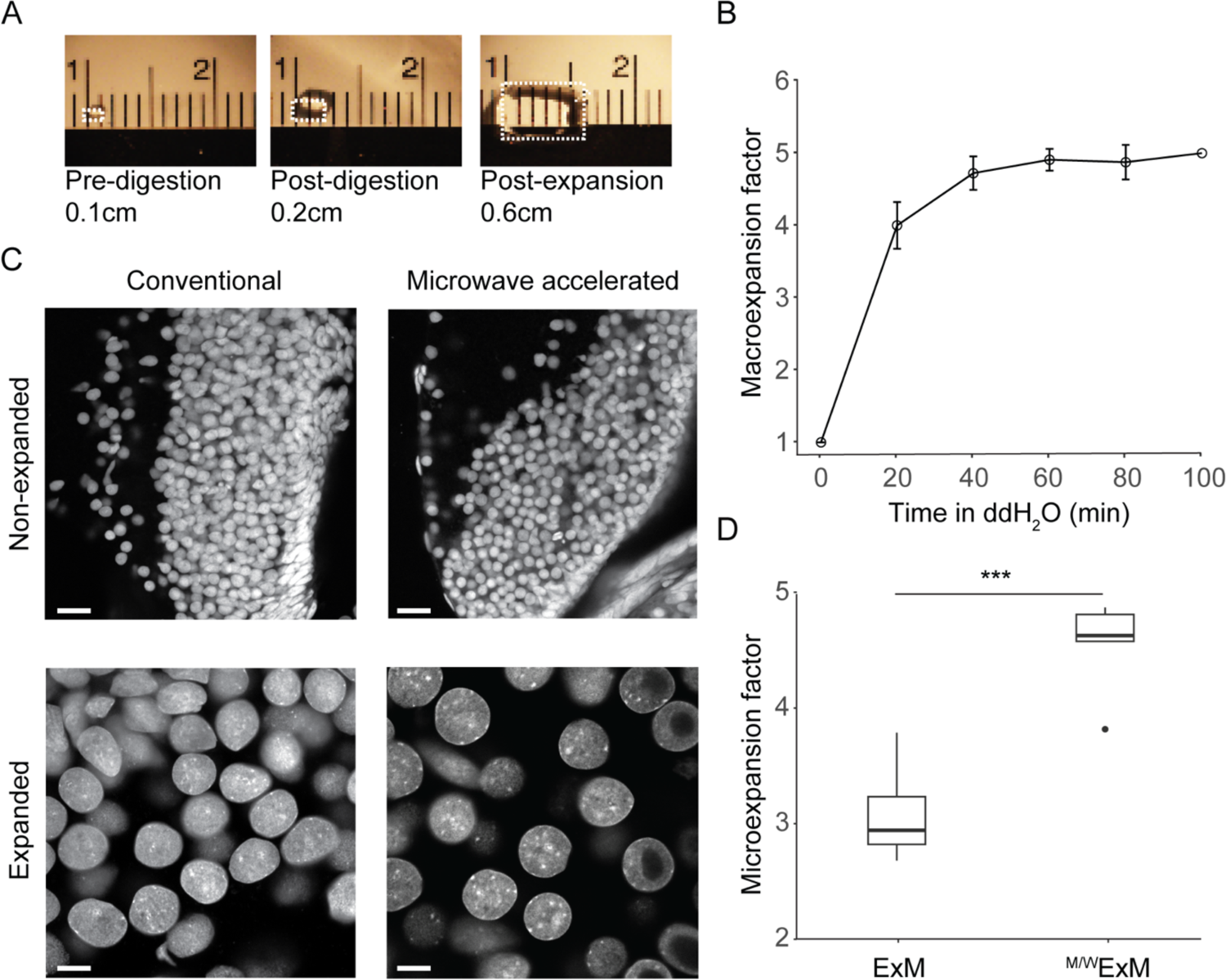
^M/W^ExM increases the magnitude of expansion. A. Representative images of a gel at different stages through the expansion protocol. Box outline depicts outline of gel at each timepoint. B. Line graph of average macroexpansion factor of the gel over time in ddH_2_O. Data shown as mean ± SEM. C. Representative images of cell nuclei in the vibratome sections of optic tectum used for the calculation of the linear microexpansion factor. Scale bar: 20μm (IHC & ^M/W^IHC); 5.7μm (ExM), 4.2μm (^M/W^ExM). D. Summary data of linear microexpansion factors achieved with conventional (ExM) and microwave-assisted (^M/W^ExM) expansion protocols. *** *p* < 0.001, independent samples t-test.

**Figure 5.**
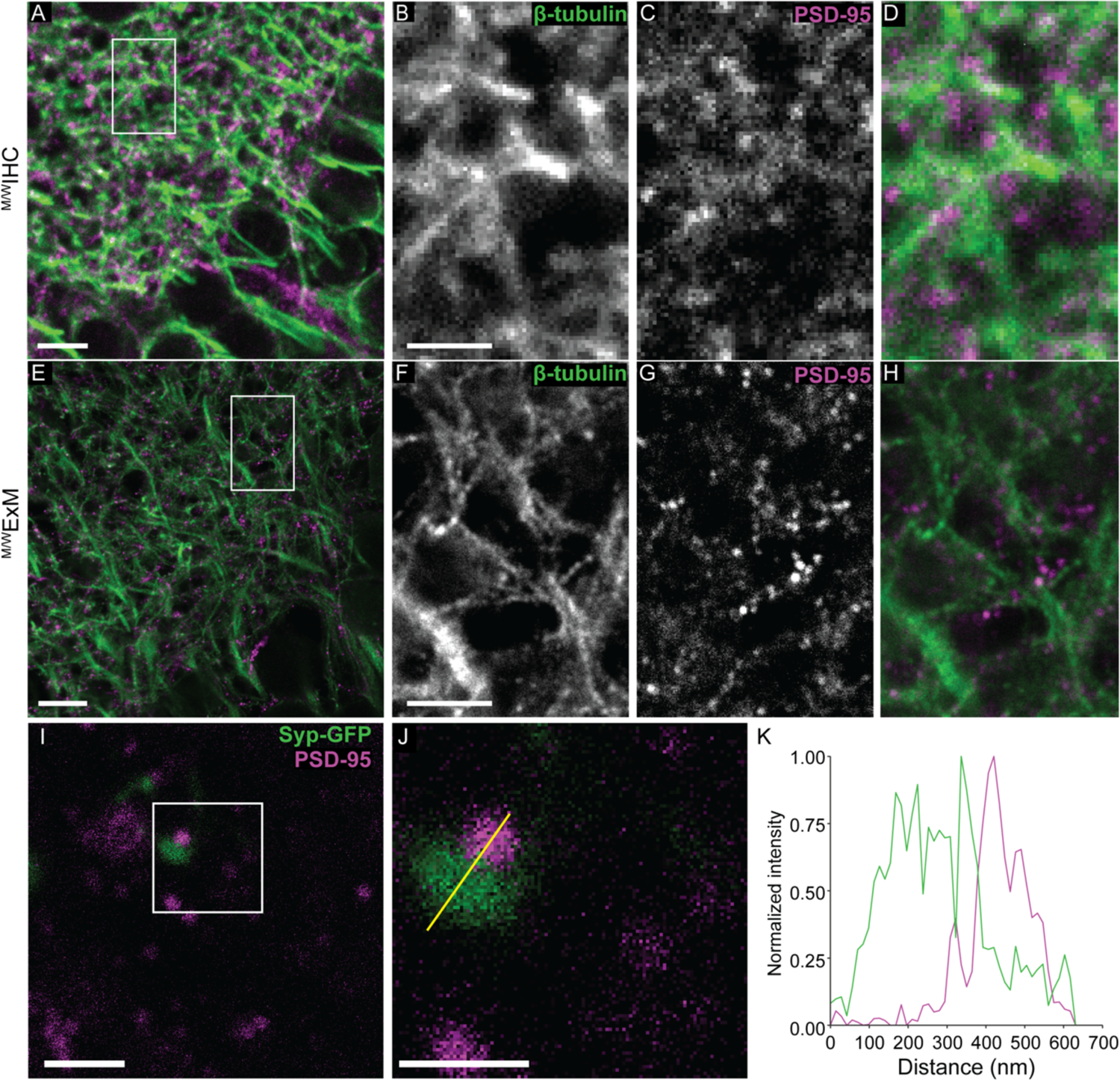
^M/W^ExM maintains the superb resolution of ExM to resolve synaptic components. A. Representative confocal image of vibratome sections of optic tectum immunostained with β-tubulin (green) and PSD-95 (magenta) using ^M/W^IHC protocol. Scale bar: 5μm. B-D. Magnification of boxed regions in (A). Scale bar: 500nm. E. Representative confocal image of vibratome sections processed with ^M/W^ExM and immunostained with β-tubulin (green) and PSD-95 (magenta). F-H. Magnification of boxed regions in (E). I. Representative confocal image of ^M/W^ExM-processed vibratome section electroporated with Synaptophysin-GFP (Syp-GFP), immunostained for GFP (green) and PSD-95 (magenta). Scale bar: 5μm. J. Magnification of the boxed region I showing a putative synapse with juxtaposing synaptophysin-GFP (green) and PSD-95 (magenta) puncta. Scale bar: 500nm. K. Fluorescence intensity plot for synaptophysin-GFP (green) and PSD-95 (magenta) for line ROI shown in J.

For *Xenopus laevis* free-floating vibratome sections, each figure panel constitutes a single optical plane or region of interest (ROI) from a single optical plane, as indicated by the figure legend (Fig. 2-5). All .tiff or .LIF files were imported and processed using built-in measurement features in ImageJ. Scale bars for expanded sections were adjusted for expansion. For each image, the linear expansion factor was determined as described below and the distance in pixels in ImageJ was adjusted to represent post-expansion size (Chen et al 2015, Tillberg et al 2016). For nuclear morphology and linear expansion analysis of SYTOX Green-stained nuclei, .tiff files were imported into Python version 3.9.13 and ROIs were calculated using the neural network algorithm Cellpose to segment cell nuclei labeled with SYTOX Green (Stringer et al 2021, Van Rossum 2009). Tiff files with ROIs identified with Cellpose were then imported into ImageJ (Schneider et al 2012). Area, circularity (*4π(area/perimeter^2^))*, and Aspect Ratio (AR; major axis/minor axis) were measured using the built-in measurement feature in ImageJ. Nuclei out of focus or in direct contact with each other (thus would result in a distorted or incomplete shape by the segmentation) were excluded from the analysis. Nuclei from progenitor cells lining the ventricle were also excluded, as their morphology and size differ from differentiated cells in the cell body layer of the optic tectum. For both measurements of nuclei morphology, the data meet the assumptions of a parametric two-way ANOVA as determined using a Shapiro-Wilkes test for normality and Levene’s test of variance equality. The area of the ROIs was used to calculate the linear microexpansion factor using this equation from Campbell et al., 2021:

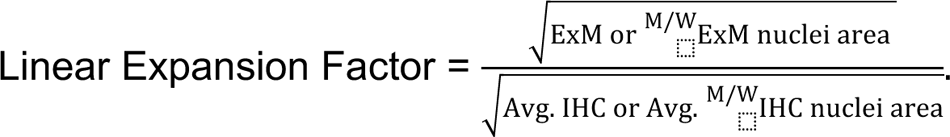

The data measuring linear expansion factor met the assumptions of an independent samples t-test as determined using a Shapiro-Wilk test for normality and Levene’s test of variance equality. For analysis of the caliber of β-tubulin fibers in Figure 2, line ROIs (5 µm) were drawn perpendicularly across discernable individual β-tubulin fibers, and the intensity along the ROI was measured using ImageJ’s Plot Profile tool. For each ROI, the minimum intensity value was subtracted as background subtraction. The full width half maximum (FWHM) value for each peak was then calculated using the findpeaks function in MATLAB (v. R2022a). These data did not meet the assumptions of a parametric one-way ANOVA as determined using a Shapiro-Wilkes test for normality and Levene’s test of variance equality, so a Kruskal-Wallis rank sum test was used. The fluorescence intensity plot in Figure 5 was done by drawing a line ROI across juxtaposing synaptophysin and PSD-95 puncta in a single optical section. The intensity across the ROI was measured using ImageJ’s Plot Profile tool. The ROI’s intensity was then normalized to its highest values. For data pertaining to *Xenopus laevis* vibratome sections, R version 4.2.1 (R Team 2023) for Mac was used for statistical analyses (Fig. 3B&D, Fig. 4E-F). Also, all graphical representations of data were done using the ggplot2 package in R version described above (v3.3.3; Fig. 3B&D, Fig. 4E-F).

For *Drosophila* whole brain samples, each figure panel constitutes a single optical plane or region of interest from a single optical plane, as indicated by the figure legend (Fig. 6-7). For data presented in Figure 6, the area under the curve analysis was performed by drawing either 20 μm (expanded) or 3.6 μm (non-expanded, calibrated by the linear microexpansion factor) line ROIs across multiple compartments of the mushroom bodies in single optical sections from a z-stack. The intensity across the ROIs was measured using ImageJ’s Plot Profile tool. Each ROI’s intensity was then normalized to its highest value. The normalized area under the curve of each individual ROI was calculated using the trapezoidal rule of integration. A two-tailed unpaired Mann-Whitney test was performed to compare the AUCs (*n* = 20-24) for a CI of 95%. *P <* 0.05 was used to determine the statistical significance for all statistical tests. Cells from both ExM and ^M/W^ExM protocols came from five different gels from at least three separate experiments. The nuclear morphology and linear expansion analysis (Fig. 7) were performed in the same way as described above. Imaging data for *Drosophila melanogaster* whole brains was analyzed and graphed in GraphPad Prism for Mac (Version 10.0.3; Fig. 7C-E).

**Figure 6.**
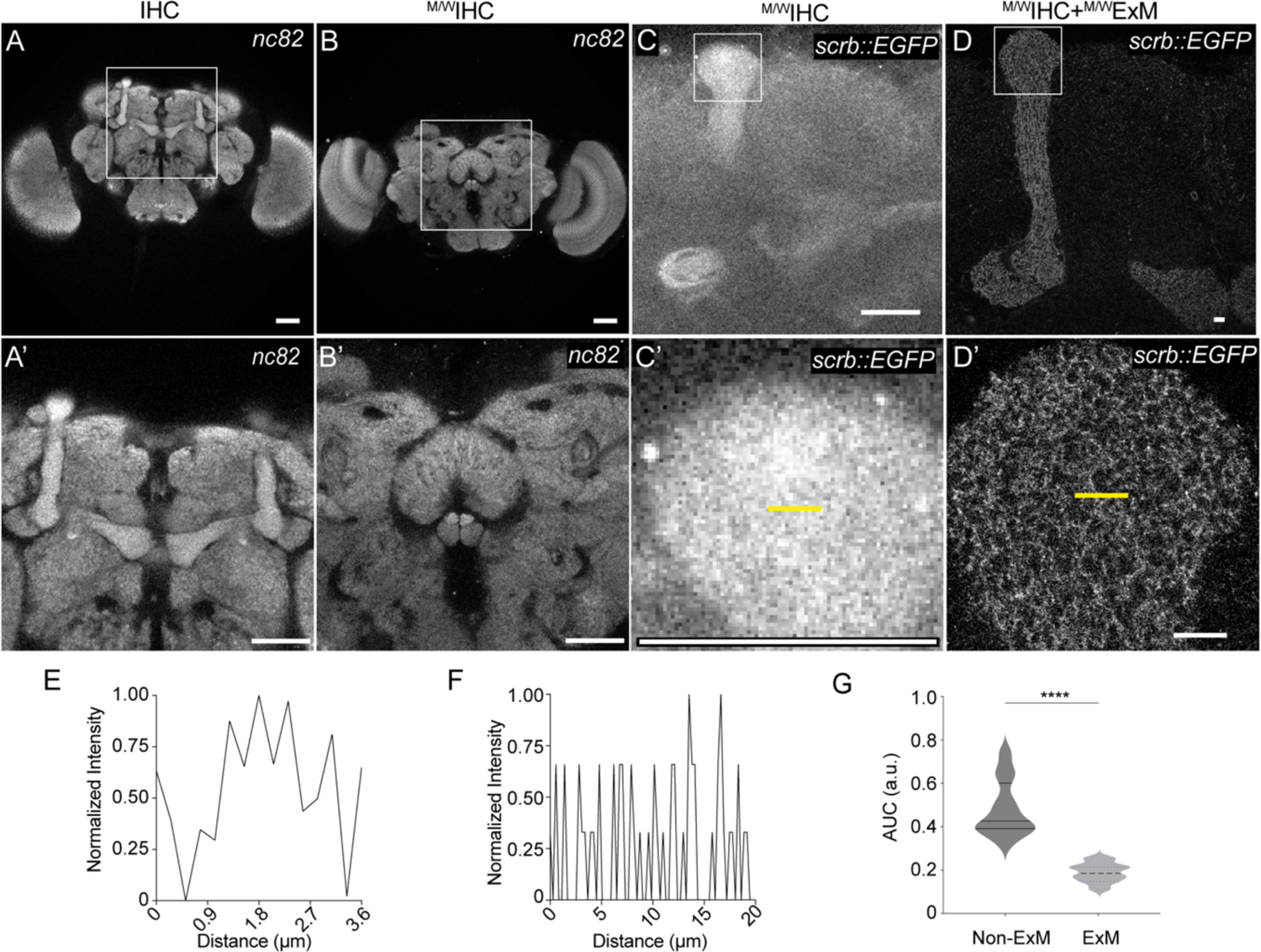
Microwave radiation accelerates immunostaining and expansion microscopy of *Drosophila* whole brain tissue. A-B, Representative single optical plane confocal images of the *Drosophila* brain immunostained with the anti-nc82 antibody using the conventional (IHC, A) and microwave-accelerated (^M/W^IHC, B) IHC protocol. C-D, Microwave-assisted immunostaining (^M/W^IHC) and expansion (^M/W^IHC-^M/W^ExM) of a Scribble-GFP-expressing *Drosophila* brain immunostained with rabbit anti-GFP. A’-D’. Magnification of boxed regions marked in A-D. A’, C’ and D’ show the *Drosophila* mushroom bodies. B’ shows the fan-shape body. Scale bars: 20 μm. E-F. Representative intensity plots of normalized intensity across line ROIs (yellow line) in ^M/W^IHC (C’) and ^M/W^IHC-^M/W^ExM (D’). G. Summary data of the area under the curve (AUC) of intensity plots from ROIs in the non-expanded (3.6 μm) and ^M/W^ExM (20μm) images. The size of the line ROI in the non-expanded images were selected to match the size of ROIs in the expanded images calibrated by the microexpansion factor. All ROIs were drawn at comparable locations within the vertical and horizontal lobes of the mushroom bodies. ****: *p* < 0.0001, Mann Whitney test, n = 20-24.

**Figure 7.**
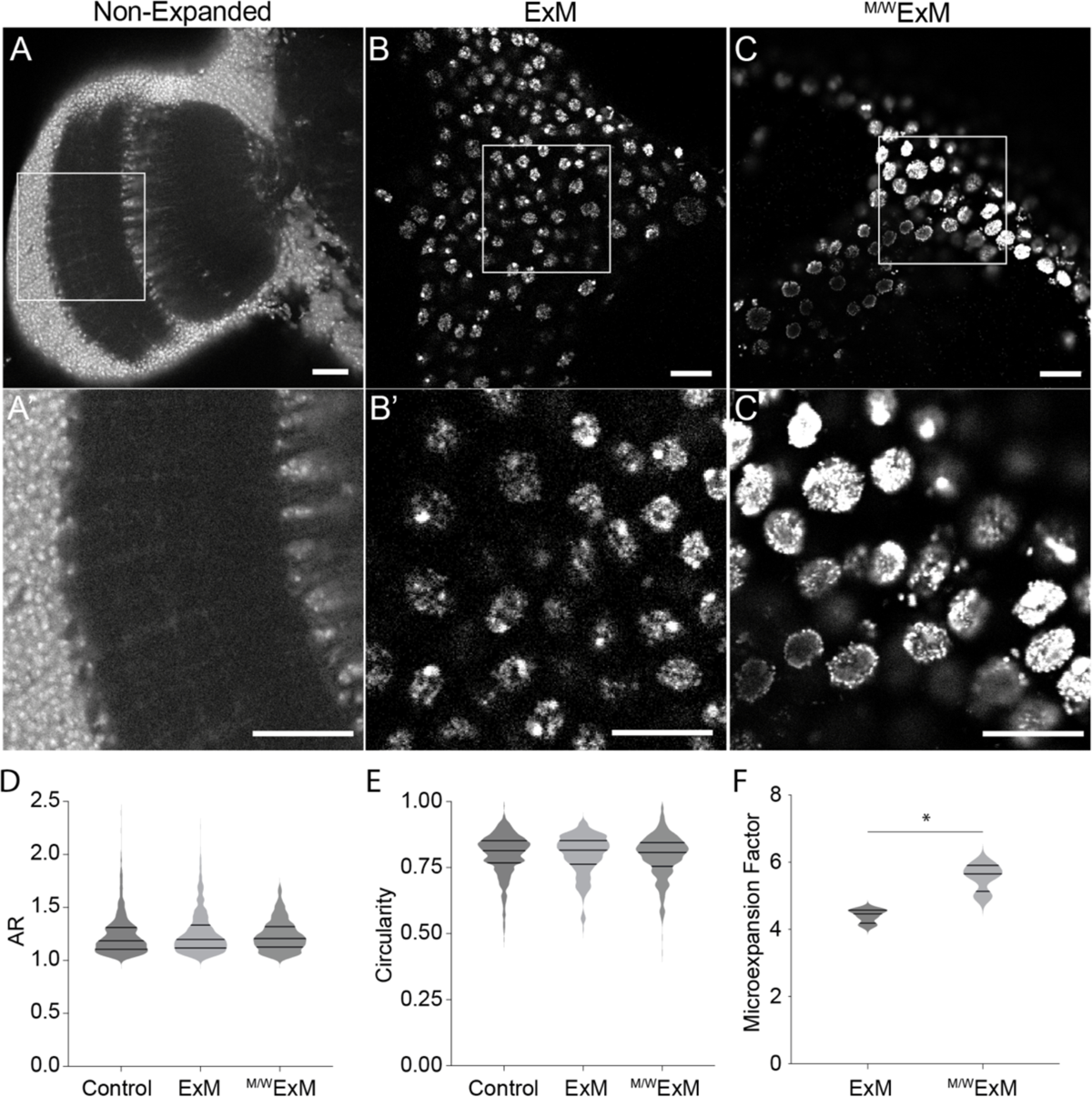
^M/W^ExM preserved nuclei morphology and significantly increased the linear expansion factor in the *Drosophila* brain tissue. A-C. Representative confocal microscopy images of the *Drosophila* optic lobe stained with SYTOX Green in (A) non-expanded, (B) ExM and (C) ^M/W^ExM samples. A’-C’. Magnification of the boxed regions in A-C. D-E. Summary data for aspect ratio (AR, D) and circularity (E) measurements of nuclei morphology, Kruskal-Wallis test; AR: *p* = 0.2079; circularity: *p* = 0.0863. F. Summary data for the linear microexpansion factor. Unpaired Mann Whitney test, *p* = 0.0286.

## 3. Results

### 3.1 Microwave radiation accelerates IHC-ExM in *Xenopus laevis* brain tissue

Neither microwave-assisted IHC nor conventional ExM had previously been tested in *Xenopus laevis*. We first tested whether the microwave-assisted IHC protocol could be adapted to fixed neural tissue of *Xenopus laevis* tadpoles. We performed immunostaining in free-floating vibratome sections of the optic tectum with anti-β-tubulin antibody, which labels microtubules in cells and has been used as a subcellular structural marker to evaluate the performance of novel immunolabeling and microscopy techniques(Bates et al 2008, Ferris et al 2009, Huang et al 2009, Tillberg et al 2016).

We adapted the ^M/W^IHC from a previously published protocol (Ferris et al 2009) and tested different key parameters (wattage, on/off duration for each cycle, number of cycles, etc.; see Table 1 for details) for the microwave processing steps. Consistent with prior reports (Bond & Kinnamon 2013, Ferris et al 2009), confocal images from sections processed with microwave-assisted IHC (^M/W^IHC Fig. 2B-B’) showed comparable immunofluorescence signal with improved signal-to-noise ratio to sections processed by the conventional IHC method (IHC, Fig. 2A-A’, also see Fig. 2F full-width at half-maximum (FWHM), mean±SEM, IHC = 0.84±0.023, N = 3 sections, n = 39 ROIs, ^M/W^IHC= 0.79±0.028, N = 3, n = 39; Z = −0.75, *p* = 0.45, Dunn’s Test for Multiple Comparisons). We then adapted an ExM protocol (Asano et al 2018) to tadpole tissue and expanded the immuno-stained vibratome sections processed by the conventional IHC method. As expected, ExM (Fig. 2C-C’) significantly improved the resolution achievable under identical IHC conditions and image acquisition parameters compared to the non-expanded section (Fig. 2A-A’). Microtubule filaments visualized by β-tubulin immunostaining could be resolved at significantly finer FWHM in ExM sections than in the non-expanded section as shown by quantified FWHM of the finest discernable individual microtubules in the images (Fig. 2F, mean±SEM: IHC = 0.84±0.023, N = 3, n = 39, ExM = 0.33±0.012 N = 3, n = 42; Z = −7.52, *p* < 0.0001, Dunn’s Test for Multiple Comparisons).

Next, we tested the feasibility of using microwave processing to accelerate the ExM protocol. There are two particularly time-consuming steps involved in the ExM protocol (Tillberg et al 2016): 1. The AcX incubation step (>6h), which facilitates the anchoring of fluorescent proteins or secondary antibody-conjugated fluorophores to the swellable gel; 2. The protease digestion step (8∼16h) homogenizes the mechanical property of the tissue and ensures it to expand isotropically. We replaced each of these steps with microwave processing (See Table 1 for details) and performed the rest of the steps as in the original adapted protocol (see method 2.9 for details). As shown in Fig 2D-D’ (^M/W^ExM), the fluorescent signals from the fluorophore-conjugated secondary antibodies persisted through the microwave processing steps. The ^M/W^ExM protocol also maintained the superior resolution of the conventional ExM protocol, as demonstrated by the microtubule structure and the quantified FWHM (Fig 2F; mean±SEM, ExM = 0.33±0.012 N = 3, n = 42, ^M/W^ExM = 0.30±0.008 N = 3, n = 54; Z = −1.28, *p* = 0.40, Dunn’s Test for Multiple Comparisons). Scales on the ExM images were registered/calibrated with the expansion factor estimated by average nuclear diameters (see next section and method for details).

### 3.2 ^M/W^ExM preserves gross nuclei morphology

The isotropic property of various expansion microscopy techniques has been well-documented (Chen et al 2015, Chozinski et al 2016, Freifeld et al 2017, Tillberg et al 2016). Isotropy of expansion has most often been validated by comparing nanoscale structures of highly stereotypical subcellular organelles such as clathrin-coated pits in images acquired with super-resolution structured illumination microscopy (SR-SIM) of the pre-expanded gels to post-expanded images acquired on regular confocal microscopes (Chen et al 2015, Chozinski et al 2016, Freifeld et al 2017, Tillberg et al 2016). Here, to verify that microwave processing does not affect the isotropic property of ExM, we compared the morphological features of cell nuclei in confocal images from sections processed by IHC, ^M/W^IHC, ExM, and ^M/W^ExM. The cell body layer of the optic tectum (OT) in *Xenopus laevis* tadpoles is composed of packed neuronal somata with relatively uniformly circular-shaped nuclei. We took advantage of this unique anatomical feature of OT and used it to estimate potential morphological distortions that may occur during expansion in the ExM and ^M/W^ExM protocols. To do this, we stained the sections with the nucleic acid stain SYTOX to visualize cell nuclei in non-expanded as well as expanded sections (Fig.3A-D). We used a neural network algorithm, Cellpose (Stringer et al 2021) to segment cell nuclei and quantified two different measurements for nuclear morphology: circularity and aspect ratio (AR) (Fig.3A’-D’). Circularity measures how closely matched the shape of a measured object is to a circle, and AR depicts the ratio of the major and minor axes. For both measurements, a value of 1 describes a perfect circle. Neither expansion (F(1) = 0.024, *p* = 0.88, two-way ANOVA) or microwave processing (F(1) = 0.352 *p* = 0.56, two-way ANOVA) had any effect on circularity (Fig. 3E). There was also no statistically significant interaction between the effects of expansion and microwave processing (mean±SEM, IHC = 0.86±0.006, N = 3 (separate sections), n = 154 (number of nuclei); ^M/W^IHC = 0.86±0.005, N = 3, n = 237; ExM = 0.85±0.007, N = 5, n = 84; ^M/W^ExM = 0.86±0.007, N = 5, n = 58; F(1) = 0.23, *p =* 0.63, two-way ANOVA) (Fig. 3E). For AR, no effect was observed for expansion (F(1) = 0.5498, *p* = 0.55, two-way ANOVA), however, a significant effect was observed for microwave processing (F(1) = 6.152, *p* = 0.029, two-way ANOVA) although the difference among different groups was extremely small (mean±SEM, IHC = 1.18±0.035, N = 3 (separate sections), n = 154 (number of nuclei); ^M/W^IHC = 1.14±0.015, N = 3, n = 237; ExM = 1.17±0.035, N = 5, n = 84; ^M/W^ExM = 1.12±0.019, N = 5, n = 58) (Fig. 3F).

As seen with circularity, there was also no statistically significant interaction between the effects of expansion and microwave processing on AR (F(1) = 0.17, *p* = 0.69, two-way ANOVA). These data are consistent with the reported isotropic property of the ExM and show that measurements of morphological properties of the nuclei may be used as an approximate evaluation for the isotropic property for different tissue processing methods. These data also suggest that the gross nuclei morphology is largely unaffected by microwave processing.

### 3.3 ^M/W^ExM increases the magnitude of the expansion

Expansion of the hydrogel is achieved by repeated 20-minute sessions of dialysis in excess volumes of H_2_O, and plateaus after several sessions (Asano et al 2018, Chen ^e^t al 201^5^). In an effort to standardize the ^M/W^ExM protocol, we took measurement of the length of gels before protease digestion and then at the beginning of each dialysis session (every 20 minutes) to determine the average time (sessions) it takes for the gel expansion to plateau in the ^M/W^ExM protocol (Fig. 4A). We then calculated the linear macroexpansion factor (Campbell et al., 2021) by normalizing the gel length at each time point to the pre-digestion length. The data showed that the gel expansion plateaued between 40 to 80 minutes after about three sessions of water dialysis (mean±SEM, maximum linear macroexpansion factor = 4.9±0.165, N = 7, n = 14, Fig. 4B).

The expansion factor of ExM can be affected by multiple factors, such as digestion duration, pH, temperature, enzyme quality, and buffer composition (Asano et al 2018, Campbell et al 2021, Park et al 2019). To ask if the ^M/W^ExM protocol maintains the expansion factor of the conventional non-microwave-assisted ExM protocol, we measured the area of individual SYTOX-stained nuclei in expanded and unexpanded sections that were processed in either conventional or microwave-assisted protocols (Fig.4C) and calculated the linear microexpansion factor (Campbell et al 2021, also see method for details). Surprisingly, the ^M/W^ExM protocol yielded a significantly higher linear expansion factor than the conventional method, as evaluated by the microexpansion factor (mean±SEM, ^M/W^ExM = 4.52±0.59, N = 5 gels, n = 65 nuclei; ExM = 3.80±0.64, N = 5, n = 86, t(138.57) = −16.57, *p* < 0.001, independent samples t-test) (Fig.4D). These data suggest that in addition to significantly reducing the total amount of the processing time of the ExM protocol, microwave radiation may also facilitate the expansion. It was also noted that the average microexpansion factor is slightly lower than the observed average macroexpansion factor, which is consistent with prior reports (Buttner et al 2021, Campbell et al 2021, Katoh 2016, Martinez et al 2020, Pernal et al 2020, Vanheusden et al 2020).

### 3.4 ^M/W^ExM maintains the superior resolution of ExM to resolve synaptic components

ExM has been demonstrated to yield superior resolution for subcellular structures in neural tissue and has since become a vital tool in neuroscience (Babi et al 2022, Eichler et al 2017, Freifeld et al 2017, Gallagher & Zhao 2021, Karagiannis & Boyden 2018, Sneve & Piatkevich 2021). To demonstrate that ^M/W^ExM retains the resolution for subcellular structures in neurons at the synaptic level as had been shown with conventional ExM, we dual-labeled free-floating vibratome tectal sections with antibodies against PSD-95 (to label excitatory synapses, Fig. 5A-B) and β-tubulin (to label the neuronal processes Fig. 5C-D). In the ^M/W^ExM image, individual PSD-95 puncta are very well resolved and much more discernable than those in the non-ExM image, suggesting much better resolution compared to the non-expanded image (Fig 5E-F). The ^M/W^ExM image also showed better signal-to-noise ratio, likely due to reduced background fluorescence and light scattering following protease digestion and water dialysis from the ExM process (Campbell et al 2021, Richardson & Lichtman 2015). To further test whether ^M/W^ExM allows the resolving of pre- and post-synaptic components of synapses, we electroporated tectal neurons with synaptophysin-GFP-expressing DNA plasmid and co-immunostained vibratome sections with transfected neurons with antibodies against GFP and PSD-95 to simultaneously label presynaptic and postsynaptic components. The presynaptic terminal (as indicated by synaptophysin-GFP) and postsynaptic terminals (as indicated by PSD-95) were clearly resolved in the ^M/W^ExM section (Fig. 5I-J), which is also shown by the normalized intensity profile drawing across juxtaposing synaptophysin/PSD-95 puncta (Fig. 5K). Together, these results demonstrate that similar to previous ExM protocols, ^M/W^ExM can be used to resolve fine subcellular structures in neural tissues such as synapses (Cahoon et al 2017, Chen et al 2015, Chozinski et al 2016, Freifeld et al 2017).

### 3.5 Microwave radiation accelerates IHC and ExM of whole brain *Drosophila* whole brain tissue

ExM of *Drosophila* tissues has been reported for embryos, larvae, and adults (Jiang et al 2018). Here, we optimized the IHC and ExM protocols for whole-mount adult *Drosophila* brains using microwave radiation. The conventional immunostaining protocol for *Drosophila* whole brain tissue takes about 6 days, excluding the fixation time (Jenett et al 2012). With microwave-assisted processing, we successfully shortened the protocol to 8 hr (Fig. 1). To test the quality of the ^M/W^IHC, we performed immunostaining with the anti-Bruchpilot antibody (nc82), which labels a key component of the active zone in drosophila melanogaster. Confocal images from ^M/W^IHC −processed brains showed dense staining in the neuropil, comparable to brains stained with the conventional IHC protocol as previously reported (Jenett et al 2012) (Fig. 6A-6B’).

Images in Figure 6 show brains stained with CF633-coupled secondary antibodies but similar results were observed using a secondary antibody with a different fluorophore (CF488, data not shown). For the ^M/W^IHC protocol, we focused on the application of microwave-assisted processing to the antibody incubation and washing steps and optimized the microwave parameters specifically for these steps. However, the fixation step may also be accelerated and enhanced using microwave radiation, as suggested by prior reports (Kahveci et al 1997).

We then adapted the microwave processing to the expansion protocol with ^M/W^IHC-processed-tissue. Again, we focused optimizing the most time-consuming anchoring and digestion steps (see Table 1 and Methods for details). For this, we stained for GFP signal in the genetically encoded Scribble::GFP protein-trap flies, *scrb::GFP^CA07683^* (Buszczak et al 2007). *Scrb* encodes a scaffolding protein known to play important roles in cell polarity determination and memory forgetting (Bilder et al 2000, Cervantes-Sandoval et al 2016). The protein trap has been reported to reveal strong expression of Scribble in two compartments of the Mushroom Bodies (MB): the α/β and γ lobes (Cervantes-Sandoval et al 2016). We detected expression patterns consistent with prior reports in the whole-mount brain tissue processed with the ^M/W^IHC-^M/W^ExM protocol (Fig. 6C-6C’). We then sought to verify that ^M/W^ExM maintains the superior resolution that is enabled by the conventional ExM methods. We compared the GFP-tagged scribble puncta in the non-expanded (Fig. 6C-6C’) and ^M/W^ExM-expanded (Fig. 6D-6D’) *scrb::GFP^CA07683^* fly brains. Confocal images of Scribble::GFP in the ^M/W^ExM brain revealed a reticular-like pattern of scribble expression in the MB neuropil (Fig. 6D-6D’), which was not discernable in the non-expanded brain (Fig. 6C-6C’).

Additionally, individual Scribble puncta are much better resolved in confocal images from ^M/W^ExM-processed brain tissue (Fig. 6D-6D’) than in the non-expanded images (Fig. 6C-6C’), which can also be seen in the intensity plots drawn across neighboring Scribble puncta (Fig. 6E-6F). Resolution is characterized as the minimum distance between two points on a sample that can be identified as distinct entities by an optical system (Li & Huang 2020). Since the separation between two points of signal will produce a decreased area under the curve (AUC) in a linear ROI, we thus quantified the area under the curve (AUC) in normalized intensity plots across multiple linear ROIs in single optical images from the ^M/W^ExM-expanded and the non-expanded brain tissue as a readout for resolution. Summary data show that AUCs from the ^M/W^ExM brain tissue are significantly lower than those from the non-expanded tissue (Fig. 6G, Mann-Whitney Test. *p <0.0001*. N = 20-24 ROIs. Median, interquartile range. IHC = 0. 4263, 0.3904-0.6007; ^M/W^ExM = 0.1897, 0.1528-0.2178), suggesting significantly increased resolution rendered by ^M/W^ExM.

To evaluate the effect of microwave radiation on the isotropic property of the ExM, we analyzed the morphological features of SYTOX-stained cell nuclei in the optic lobes of the *Drosophila* brain using the same protocol as described in 3.2. We compared the nuclei morphology in non-expanded brains (Fig. 7A-A’) and brains expanded with conventional ExM (Fig. 7B-B’), as well as brains expanded with ^M/W^ExM (Fig. 7C-C’). We found no statistically significant differences among the three groups for each of the morphology descriptors measured (Fig. 7D-E, Aspect Ratio: Kruskal-Wallis Test. H (2) = 3.142. *p* = 0.2079. Median, interquartile range. Control = 1.186, 1.107-1.312, N = 4 samples, n = 677 number of nuclei; ExM = 1.2, 1.119-1.333, N = 3, n = 819; ^M/W^ExM = 1.207, 1.126 −1.319, N = 3, n = 782; Circularity: Kruskal-Wallis Test. H (2) = 4.901 *p* = 0.0863. Median, interquartile range. Control = 0.814, 0.768-0.852, N = 4, n = 940; ExM = 0.816, 0.763-0.853, N = 3, n = 643; ^M/W^ExM = 0.807, 0.7545-0.845, N = 3, n = 629). To test if the ^M/W^ExM protocol maintains the expansion factor of the conventional ExM protocol, we also calculated the linear microexpansion factor as described above in 3.3. The ^M/W^ExM treatment also yielded significantly higher microexpansion factors than conventional ExM in *Drosophila* brain tissue tadpoles (Fig. 7F, median, interquartile range, ^M/W^ExM = 5.652, 5.133-5.903. N = 4 (gels), n = 352 (number of nuclei); ExM = 4.459, 4.183-4.571. N = 4 (gels), n = 352 (number of nuclei); U = 0, *p =* 0.0286, Mann-Whitney test), consistent with our observation in the brain tissue of *Xenopus laevis*. These results demonstrate that ^M/W^ExM successfully accelerates the conventional IHC and ExM protocols without compromising tissue integrity and expansion efficiency.

## 4. Discussion

With its easy accessibility and wide applicability, ExM has been enthusiastically embraced by the community of biological research since its debut (Chen et al 2015). It has been adapted to many different species as well as different tissue types (Alonso 2022, Deshpande et al 2017, Freifeld et al 2017, Gaudreau-Lapierre et al 2021, Gorilak et al 2021, Gotz et al 2020, Jiang et al 2018, Kao & Nodine 2021, Kong & Loncarek 2021, Yu et al 2020). Here we successfully adapted the ExM method to the brain tissue of *Xenopus laevis* tadpoles for the first time and demonstrated that a 3∼4x linear expansion can be achieved with the conventional ExM protocol, without compromising tissue integrity and morphology of subcellular structures. More importantly, by applying microwave-assisted processing to the most time-consuming steps in the ExM protocol, we demonstrated that the protocol can be significantly accelerated. Hence it greatly increases the experimental efficiency. The ^M/W^ExM protocol maintains the superior resolution and signal-to-noise ratio rendered by the original ExM protocol and yields significantly higher linear expansion with the same reagents and steps. The microwave-assisted acceleration of the ExM protocol can be readily adapted to other species, as we have demonstrated with the whole-mount brain tissue of *Drosophila melanogaster*. In addition to the increased efficiency, with precisely controlled parameters of each step, the microwave-assisted protocol also helps to improve the consistency of the outcome of the protocol.

In general, several parameters need to be considered when applying microwave radiation to accelerate either the immunostaining or expansion processes. To optimize the outcome (e.g. time, expansion efficacy, signal-to-noise ratio, preservation of morphology) while avoiding overheating, it is important to test for different power levels (wattage), the duration and number of cycles, and intervals between cycles, for any given the volume of fluid and tissue type. For reference, we listed the optimized parameters that we found with different applications presented here (Table 1). Once the power, duration and interval are set, the number of M/W cycles for each step can be a convenient variable to test. For example, for the digestion step, we found that too many cycles led to degradation of the tissue and a decrease in the fluorescence signal. Too few cycles, on the other hand, compromised the efficiency of digestion which led to increased background noise and decreased magnitude of expansion. This is consistent with our observation that M/W facilitates the expansion process and leads to significantly increased linear expansion. We also found that there is not a linear relationship between the length of incubation duration used in the conventional protocol and the number of M/W cycles used to substitute it, i.e. each step (e.g. AcX treatment, digestion, antibody incubation) must be determined independently of the conventional protocol.

For the ^M/W^IHC protocol, as for the conventional IHC protocols, the choice of M/W parameters can vary by specific primary antibodies, the preparation of tissues, and different types of tissues, so must be independently determined for each new protocol. As previously reported, the concentration of the primary and secondary antibodies may need to be increased when adapting to ^M/W^IHC or ExM protocols, especially for ExM protocols due to reduction of fluorescence by digestion and expansion (Campbell et al 2021, Ferris et al 2009, Tillberg et al 2016). However, no extra increase in antibody concentration is needed for the ^M/W^IHC-^M/W^ExM protocol compared to the conventional IHC-ExM protocol. In addition, we found that the previously reported need for increased antibody concentration maybe ameliorated by increasing the number of M/W cycles. For example, we found that increasing the number of M/W cycles from 3 to 7 for PSD-95 antibody incubation (with the same M/W power, duration and interval parameters) significantly increased the antibody penetration and binding, and allowed us to achieve better labeling results with the same concentrations of primary antibody as used for conventional IHC (see Methods for details). Another factor to pay attention to is the number of primary antibodies used for co-immunostaining. We consistently observed that when working with multiple antibodies, the total number of M/W cycles must be increased to achieve the same level of antibody penetration and labeling (Fig.5). Increasing the microwave wattage for fewer cycles may be possible, but it should be combined with increased interval (off-cycle) duration to allow more time for the tissue (and solution) to cool down between cycles.

ExM is a powerful tool to overcome traditional optical diffraction limits in immunolabeled specimens. The microwave-assisted protocol greatly increases the efficiency of the expansion protocol, without compromising the tissue integrity, while maintaining the superior resolution enabled by the conventional expansion method. Many variations of the expansion microscopy methods have been developed so far. These alternative methods allowed for either a significantly higher level of expansion (Chang et al 2017) or better immunolabeling (Mantyla et al 2023). It is possible that the ^M/W^ExM protocol may also be adapted to these expansion microscopy protocols.

## CrediT Authorship contribution statement

**M.R.B.:** Conceptualization, Methodology, Formal Analysis, Investigation, Visualization, Writing. **J.C.M-C.:** Methodology, Formal Analysis, Investigation, Visualization, Writing. **N.B.Q.:** Methodology, Writing. **A.J.:** Investigation, Formal Analysis**. D.A.A.:** Methodology. **J.M.L.:** Formal Analysis. **E.I.:** Methodology, Investigation. **T.M.S.:** Methodology, Investigation. **I.C-S.:** Methodology, Visualization, Supervision, Writing. **H-y.H.:** Conceptualization, Methodology, Funding acquisition, Visualization, Supervision, Writing.

## Acknowledgements

We thank Reshmi Bera for her help in preparation of reagents. We also thank the He lab for their feedback and input into the preparation of this manuscript. This work was supported by the NIH (KL2TR001432, R01NS133441 to H-Y H) and GU (AY2021 Research Infrastructure Award).

## Data availability

Data will be made available upon request.

